# Differentiation of mouse embryonic stem cells into cells with spermatogonia-like morphology with chemical intervention-dependent increased gene expression of LIM homeobox 1 (*Lhx1*)

**DOI:** 10.1101/2021.10.12.464060

**Authors:** Cameron Moshfegh, Sebastian G. Rambow, Seraina A. Domenig, Aldona Pieńkowska-Schelling, Ulrich Bleul, Viola Vogel

## Abstract

During the development of the male germline, spermatogonial stem cells (SSCs) originate from gonocytes that differentiate from primordial germ cells (PGCs). In the developing and regenerating mouse testis, expression of the gene LIM homeobox 1 (*Lhx1*) marks the most undifferentiated SSCs. However, an enrichment of *Lhx1* expression in spermatogonia-like cells generated *in vitro* has not been reported so far. Previously, it was shown that a chemical intervention in male mouse embryonic stem (ES) cells in serum culture, including a timed combination of the SIRT1 inhibitor Ex-527, the DNA methyltransferase inhibitor RG-108 and the electrophilic redox cycling compound tert-butylhydroquinone (tBHQ), was associated with molecular markers of the PGC to gonocyte differentiation process. Here, we report the *in vitro* differentiation of male mouse ES cells, cultured under dual chemical inhibition of GSK3β and MEK (termed 2i) with leukemia inhibitory factor (LIF) (termed 2iL) and serum, into cells with spermatogonia-like morphology (CSMs) and population-averaged expression of spermatogonia-specific genes. This was achieved by the removal of 2iL and a specific schedule of 2 partial medium replacements per day with alternating 8-hour and 16-hour intervals over a period of 32 days. Combination of this new cell culture protocol with the previously reported chemical intervention in ES cells changed the population-averaged expression of spermatogonia- and gonocyte-specific genes during the differentiation process and increased the population-averaged gene expression of *Lhx1* in the resulting CSMs compared to CSMs without chemical intervention. Furthermore, we detected single CSMs with a strong nuclear LHX1/5 protein signal only in the chemical intervention group. Our results provide the first experimental evidence for the generation of CSMs with an enrichment of *Lhx1* expression *in vitro*. We propose that further investigation of the CSMs generated with this *in vitro* system may provide new insights into male germline and stem cell development.

## Introduction

Embryonic stem (ES) cells are pluripotent stem (PS) cells determined by their potential to differentiate into the germline and all somatic lineages [1]. They can function as a cell model to investigate developmental processes *in vitro*, such as the development of the male germline [2]. Primordial germ cells (PGCs), the developmentally first cells of the germline, are induced from epiblast cells on around embryonic day (E)6.25 by external signals, consisting of bone morphogenetic protein 4 (BMP4) signaling [2, 3]. Gonocytes are induced from PGCs on around E12.5-E13.5 [2-4]. The stem cells of the male germline, defined as spermatogonial stem cells (SSCs), originate on around postnatal day (P)3-P6 from gonocytes [5]. Previous efforts to reconstitute this sequence of developmental stages *in vitro* first induced ES cells into epiblast-like cells, which in turn were induced into PGC-like cells [6, 7], and then further differentiated into spermatogonia-like cells [8]. PGC-like cells and spermatogonia-like cells both had the capacity to generate offspring after transplantation into testes of germ cell-deficient neonatal mice [6-8]. However, spermatogonia-like cells lacked enriched expression of important SSC marker genes such as LIM homeobox 1 (*Lhx1*) [8], which is a marker gene for the most undifferentiated SSCs in the developing and regenerating mouse testis [9, 10]. This observation suggests that some important aspects of spermatogonia development have not yet been reconstituted *in vitro*. Dual chemical inhibition of GSK3β and MEK (termed 2i) with leukemia inhibitory factor (LIF) (termed 2iL), is routinely used *in vitro* to stabilize the so-called ground state of pluripotency in ES cells, corresponding to cells of the inner cell mass (ICM) of the blastocyst [1]. However, it was reported that mouse ES cells *in vitro*, depending on culture conditions, exist within a spectrum of pluripotent states that influences their commitment towards the germline or somatic lineages [11]. 2i induced a PGC-like state in ES cells, while LIF inhibited this PGC-like state [11]. 2i also increased the expression of DAZL in ES cells, an RNA-binding protein and marker of late PGCs which is expressed only in a subset of cells of the ICM, while serum increased heterogeneity of pluripotent states [12]. Furthermore, it was shown for male mouse ES cells cultured in serum that a chemical intervention, including a timed combination of the SIRT1 inhibitor Ex-527, the DNA methyltransferase inhibitor RG-108 and the electrophilic redox cycling compound tert-butylhydroquinone (tBHQ), was associated with molecular markers of the PGC to gonocyte differentiation process [13]. The aim of this study was to investigate whether the combination of a new cell culture protocol based on 2iL and serum with the previously reported chemical intervention [13] can induce aspects of spermatogonia development in male mouse ES cells *in vitro*. We found that male mouse ES cells cultured in 2iL and serum could be differentiated into cells with spermatogonia-like morphology (CSMs) by removal of 2iL and a specific schedule of partial medium replacements, and that a combination of this new cell culture protocol with the previously reported chemical intervention resulted in the first observation of CSMs *in vitro* with enriched *Lhx1* expression. Further investigation of these cells may provide new insights into male germline and stem cell development.

## Results and Discussion

To investigate aspects of spermatogonia development in male mouse ES cells, we decided to use 2iL and serum as the basis for the culture, based on previous reports that serum permits a broader spectrum of pluripotent states and downstream lineages [11], that 2i induces a PGC-like state in ES cells and facilitates downstream PGC-like differentiation from ES cells [11, 12], and that 2iL supports the ground state of pluripotency [1]. Using this 2iL and serum culture basis, we designed a combined cell culture protocol without (control) and with chemical intervention (intervention) (Fig 1), based on a previous report that the chemical intervention was associated with the induction of molecular markers of the PGC to gonocyte differentiation process in male mouse ES cells in serum culture [13].

**Fig 1.**
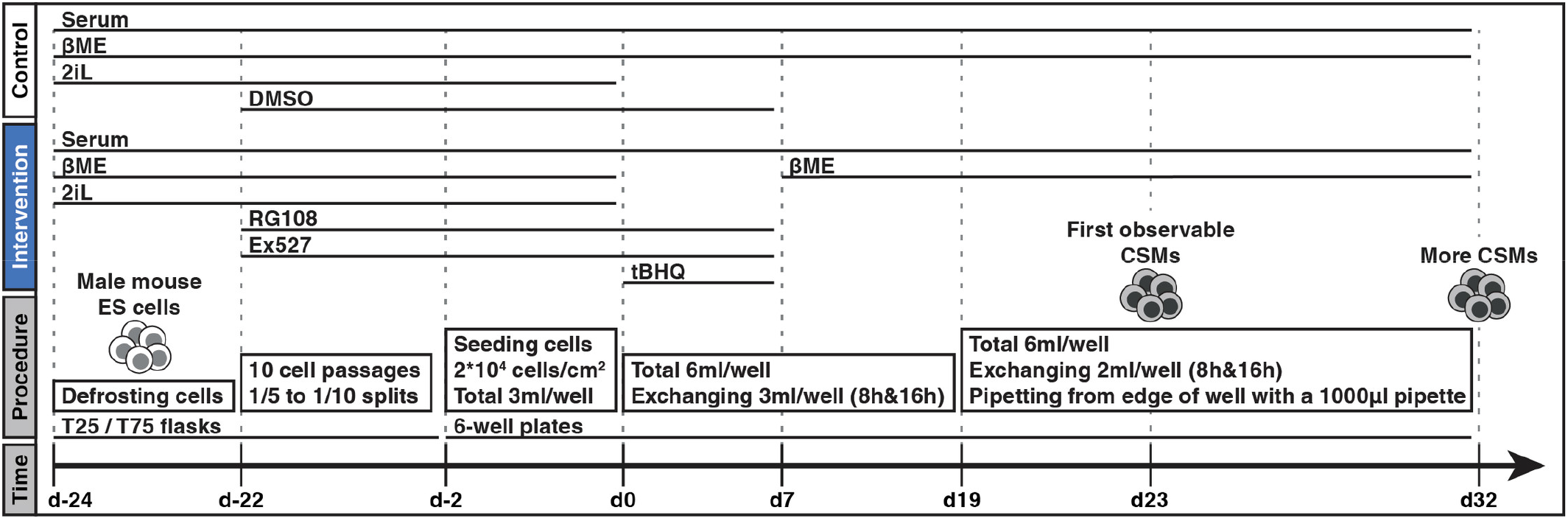
Schematic procedure for the differentiation of male mouse ES cells into CSMs. A schematic description of the procedure in the control (white) and intervention group (blue) for the differentiation of male mouse ES cells (day -24) into CSMs (day 32, first observable CSMs on day 23). See Materials and Methods for a detailed description.

The intervention consisted of a sequence of different culture conditions. After defrosting on day -24 of the protocol, male mouse ES cells were cultured for 10 cell passages in 2iL and serum with the SIRT1 inhibitor Ex-527 and the DNA methyltransferase inhibitor RG-108 in the presence of β-mercaptoethanol (βME). The cells were then seeded into 6-well plates and cultured for 2 days under the same culture conditions until day 0. For the next 7 days, the electrophilic redox cycling compound tBHQ was added and 2iL and βME were removed, while the medium was partially replaced twice daily from this point onwards (half of volume at hour 8 and hour 16). On day 7, cells were cultured for another 25 days in serum and βME without 2iL and the other chemicals. From day 19 onwards, the volume of partial medium replacements was changed to one third of the well volume, while keeping the same replacement frequency. The control group always contained βME and chemicals were replaced with dimethyl sulfoxide (DMSO) at the same final concentration, while all other parameters were kept the same as in the intervention group (see Materials and Methods and Fig 1).

We purchased male mouse C57BL/6N ES cells from Millipore and confirmed their male genetic sex (xy) by fluorescence in situ hybridization (FISH) (S1 Fig). We also found aneuploidy in these ES cells with a duplication of chromosome 8 in all three analyzed metaphase spreads (S1 Fig). Trisomy 8 is a common chromosomal abnormality in ES cells cultured long-term *in vitro*, as it confers a selective proliferation advantage [14]. Although ES cells with trisomy 8 have a diminished efficiency to contribute to the germline of chimeric mice [14], a diminished *in vitro* germline developmental potential for ES cells with duplication of chromosome 8 has not been reported. Therefore, we continued to use these ES cells.

Typical for ES cells [6], on day 0, the culture consisted of compact multicellular colonies, while on day 7 it had grown to confluent multilayered cell sheets of variable thickness with some visible debris from dead cells in both groups (Fig 2). From day 0 to day 7, the continued presence of βME supported cell viability of the control group, while tBHQ-induced nuclear factor erythroid 2-related factor 2 (NRF2) signaling rescued cell viability of the intervention group in the absence of βME as previously shown [13]. Thicker cell sheets with holes in some areas and reduced debris were observable on day 19 in both groups. Starting on day 23 in both groups, these cell sheets progressively dissociated into round mostly non-adherent cells, described here as CSMs due to their morphological similarity to previously reported isolated SSCs from P4.5-P7.5 mouse testis [15, 16]. By day 32 a large portion of the cell sheets had dissociated into CSMs in both groups (Fig 2). This process progressed faster and more completely in the intervention group than in the control group (S2 Fig). This may suggest a faster or changed differentiation process in the intervention group compared to the control group. CSMs were morphologically indistinguishable between both groups. The appearance of CSMs in the control group is the first reported observation that male mouse ES cells in 2iL and serum culture can differentiate into CSMs by removal of 2iL without additional exogenous biological or chemical factors. However, we noticed that the twice daily partial medium replacement at hour 8 and hour 16 with a third of the volume from day 19 onwards was necessary for the appearance of CSMs. When we replaced the medium twice daily every 12 hours with half of the volume, the cell sheets remained intact and CSMs did not appear in either group. This observation suggests that endogenously produced soluble factors may be necessary for the transition of cell sheets into CSMs. Interestingly, it was previously reported that the transition of gonocytes into spermatogonia in the mouse required FGF signaling [17] and that accumulation of endogenously produced FGF2 in the culture medium was essential for the survival and proliferation of cultured mouse SSCs [18]. However, we did not further investigate soluble factors. We then asked if the population-averaged expression of genes specific to spermatogonia development supported the view that the dissociation of cell sheets into CSMs showed aspects of the transition of gonocytes into spermatogonia, and how the gene expression, particularly in the resulting CSMs, was different between the intervention and control group. For this, we collected cells on day 0, day 7, day 19 and day 32 for real-time PCR analysis to detect mRNA levels. From day 0 to day 19, all cells were adherent and collected. On day 32, because our focus was on the CSMs, we only collected non-adherent cells by gentle rinsing, capturing most of the CSMs. The gene expression of the adherent cells on day 32 was not analyzed.

**Fig 2.**
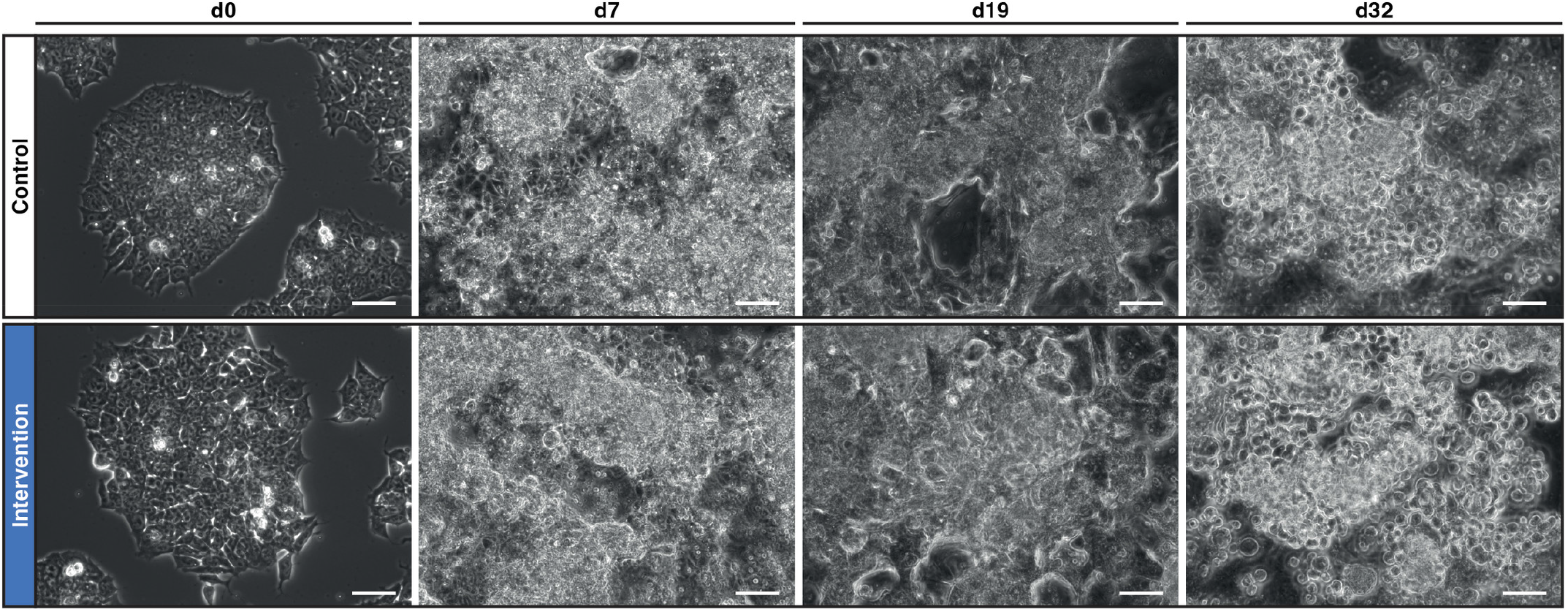
Phase contrast images during the differentiation process. Phase contrast images during the differentiation process of ES cells to CSMs in the control (white) and intervention group (blue) on day 0, day 7, day 19 and day 32. On day 0, ES cells grew as adherent compact multicellular colonies. On day 7, cells had grown to confluent multilayered cell sheets of variable thickness with some visible debris from dead cells in both groups. On day 19, the cell sheets had grown thicker with holes in some areas and reduced debris in both groups. On day 32, progressive cellular dissociation of the cell sheets had resulted in CSMs that appeared as round mostly non-adherent cells in both groups. This process was first observable on day 23 in both groups but progressed faster and more completely in the intervention group than in the control group (S2 Fig). CSMs were morphologically indistinguishable between both groups. Scale bar = 50 μm.

We found that expression of *Zbtb16, Gfra1, Bmi1, Kit* and *Klf4* was most upregulated on day 32 (containing CSMs) relative to all other analyzed time points in both groups (Fig 3). *Zbtb16* and *Gfra1* are marker genes of SSCs but are already expressed in some gonocytes from E18.5-P1.5 onwards [17]. *Bmi1* is a marker gene of SSCs [19] but is also expressed in gonocytes from E16.5 onwards [20]. *Kit* is a marker gene of differentiating spermatogonia from P2-P6 onwards [21-23], including the first round of spermatogenesis that begins directly from gonocytes and omits the SSC-stage [5]. *Klf4* is a pluripotency-associated gene [24, 25] but is also expressed in SSCs [26], differentiating spermatogonia and spermatids [20]. Therefore, these results suggest that aspects of spermatogonia development and differentiation were present on day 32 in both groups.

**Fig 3.**
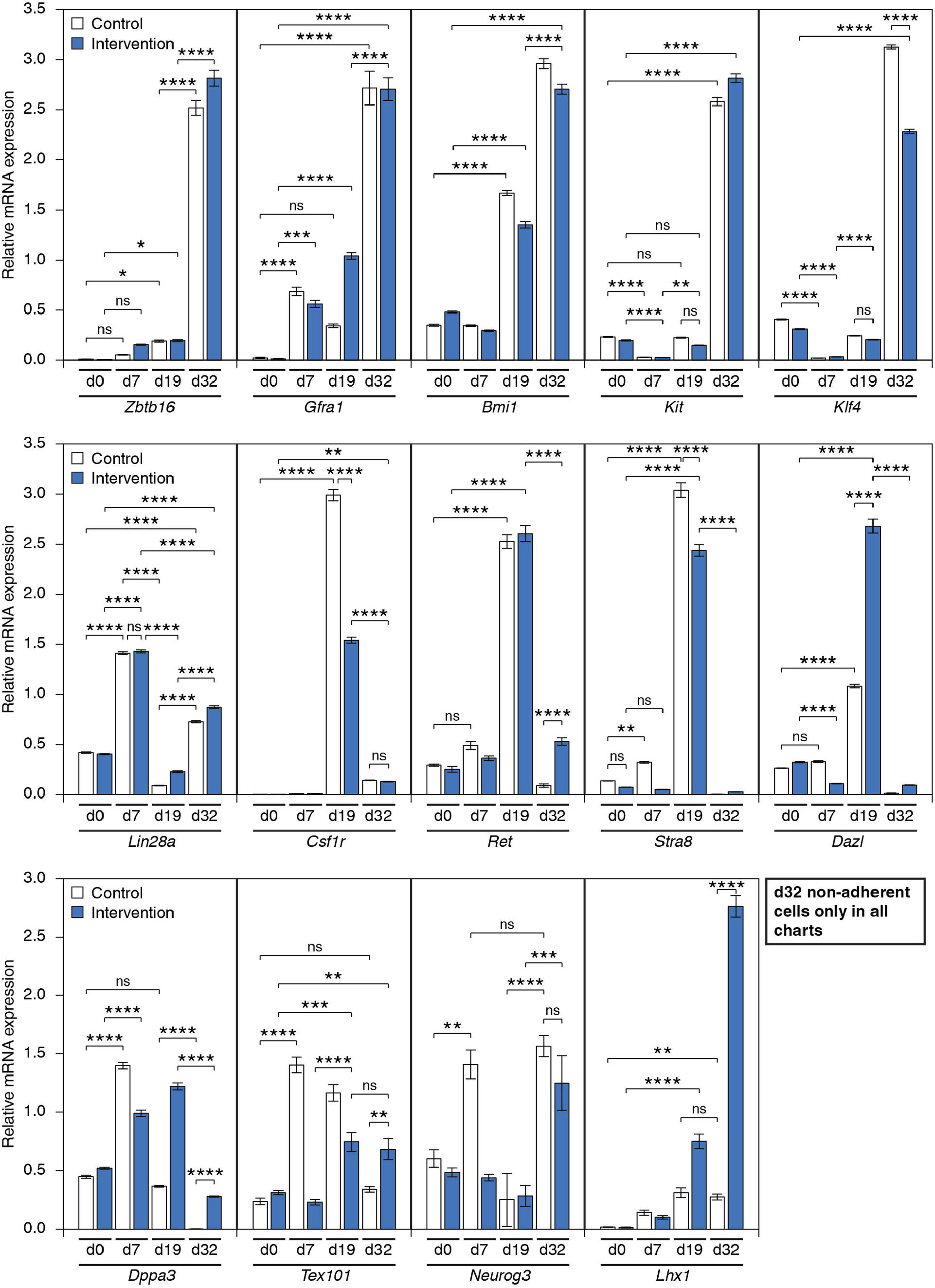
Expression of genes specific to spermatogonia development during the differentiation process. Real-time PCR analysis of genes specific to spermatogonia development during the differentiation process of ES cells to CSMs in the control (white) and intervention group (blue) on day 0, day 7, day 19 and day 32. From day 0 to day 19, all cells were adherent and collected, while on day 32, only non-adherent cells were collected. Expression of *Zbtb16, Gfra1, Bmi1, Kit* and *Klf4* was most upregulated on day 32 relative to day 0 in both groups. Expression of *Lin28a* was upregulated on day 32 and day 7 relative to day 0 in both groups. Expression of *Csf1r, Ret, Stra8* and *Dazl* showed the highest upregulation on day 19 in both groups, while *Csf1r* expression was also upregulated on day 32 relative to day 0 in both groups. Expression of *Dppa3, Tex101, Neurog3* and *Lhx1* was differentially regulated depending on the chemical intervention. Expression of *Lhx1* on day 32 was upregulated relative to day 0 in both groups and increased in the intervention group compared to the control group. Relative gene expression levels were calculated based on the ΔΔCt method with normalization using *Gapdh*. Error bars indicate standard error of the mean (SEM). * p<0.05, ** p<0.01, *** p<0.001, **** p<0.0001, ns non-significant. PCR reactions were performed with technical replicates of 5 (n=5) from pooled samples.

Furthermore, *Lin28a* expression was upregulated on day 32 relative to day 0 but showed the highest upregulation on day 7 in both groups. *Lin28a* is a marker gene of SSCs [27] but is also expressed in PGCs on E13.5 [28]. Expression of *Csf1r, Ret, Stra8* and *Dazl* showed the highest upregulation on day 19 in both groups, while *Csf1r* expression was also upregulated on day 32 relative to day 0 in both groups (Fig 3). *Csf1r* is expressed in SSCs on P6 [29]. *Ret* is expressed together with *Gfra1* and *Zbtb16* in SSCs as well as in some gonocytes from E18.5-P1.5 onwards, while *Stra8* is expressed in a subset of these gonocytes [17]. DAZL mediates a broad translational program to regulate proliferation and differentiation in the germline and is robustly expressed in male gonocytes on P0 and P4 [30]. Therefore, these results further suggest that aspects of gonocyte development preceded the transition of cell sheets into CSMs.

We then found differences in the expression of *Dppa3* and *Tex101* in the intervention group compared to the control group (Fig 3). *Dppa3* is expressed in PGCs and in male gonocytes until E15.5 [31, 32], and then its expression gradually decreases until it is no longer detectable on P1 or later [31]. *Tex101* is expressed in early gonocytes from E14-E16 onwards until early spermatogonia on P8-P10, in spermatocytes and spermatids, but not in PGCs, during gonocyte induction on E12.5-E13.5 or in spermatogonia from P12 onwards [33]. *Dppa3* expression was upregulated on day 7 relative to day 0 in both groups. In the intervention group, *Dppa3* expression remained at this level on day 19 and was downregulated to a level like day 0 on day 32. In the control group, *Dppa3* expression was downregulated already on day 19 to a level like day 0 and was further downregulated around 160-fold on day 32. In the intervention group, *Tex101* expression was upregulated on day 19 relative to day 0 but not earlier and remained at that level on day 32. This means that *Tex101* expression was not upregulated during the chemical intervention. In the control group, *Tex101* expression was upregulated already on day 7 relative to day 0, remained at that level on day 19 and was downregulated on day 32. This suggests that some aspects of spermatogonia development up to P1 and P8 progressed later in the intervention group compared to the control group.

We found further differences in the expression of *Neurog3* and *Lhx1* in the intervention group compared to the control group that suggested a changed differentiation process. *Neurog3* is expressed in undifferentiated spermatogonia from P3-P6 onwards but not in gonocytes, differentiating spermatogonia or other germ cells [5, 34]. Although not all adult SSCs express *Neurog3* [27, 35], all adult spermatogenic cells originate from these *Neurog3*-expressing spermatogonia established after birth [5, 34]. Therefore, upregulation of *Neurog3* expression was expected only when CSMs were present (day 32) but not otherwise. *Lhx1* is a marker gene of the most undifferentiated SSCs in the developing (P7-P10, potentially earlier) and regenerating mouse testis [9, 10]. In the intervention group, *Neurog3* expression was only upregulated on day 32 relative to day 0 (Fig 3). In the control group, *Neurog3* expression was upregulated only on day 7 and day 32 relative to day 0 (Fig 3). The expected upregulation of *Neurog3* expression only when CSMs were present (day 32) was confirmed in the intervention group but not in the control group. *Lhx1* expression was upregulated on day 32 relative to day 0 in both groups. However, the expression of *Lhx1* on day 32 was around 10-fold higher in the intervention group compared to the control group. Among the analyzed genes with an upregulated expression on day 32 relative to the other analyzed time points *Lhx1* showed the largest difference between the two groups. This suggests that the differentiation process in the intervention group compared to the control group was fundamentally different, leading to an increased population-averaged expression of a marker gene of the most undifferentiated SSCs (*Lhx1*) in the resulting CSMs in the intervention group.

We then tested for LHX1 protein in single cells on day 32 compared to mouse embryonic fibroblasts (MEFs) and ES cells by immunofluorescence using an antibody that detects LHX1/5 (Developmental Studies Hybridoma Bank, 4F2). Congruence of the LHX1/5 and DAPI signal in the images and z-axis profiles defined a nuclear signal. We found that LHX1/5 signal was detectable in the cytoplasm and nucleus of some cells in all analyzed samples. A strong nuclear LHX1/5 signal was detected in 3 out of 187 cells in CSMs in the intervention group but not in the other analyzed cells (MEFs: 160 cells, ES cells: 226 cells, CSMs in control group: 144 cells) (Fig 4; S3 and S4 Fig). As LHX1 is a transcription factor and shows a strong nuclear signal in SSCs from P7 mouse testis [10], a strong nuclear LHX1/5 signal supports this function. The small fraction of CSMs with a strong nuclear LHX1/5 signal may result from a combination of increased *Lhx1* expression in only a small fraction of CSMs, post-transcriptional, and post-translational regulation. These results support the population-averaged gene expression analysis of *Lhx1*. This represents the first experimental evidence for the *in vitro* generation of CSMs with an enrichment of *Lhx1* expression. Our results identify genes involved in spermatogonia development, including *Dppa3, Tex101, Neurog3* and *Lhx1*, as differentially expressed during early and late stages of the *in vitro* differentiation process, depending on a chemical intervention. How the chemical intervention achieves this was not studied here and may be a subject of future experiments. Further studies based on this cellular *in vitro* system may provide insights into new aspects of male germline and stem cell development.

**Fig 4.**
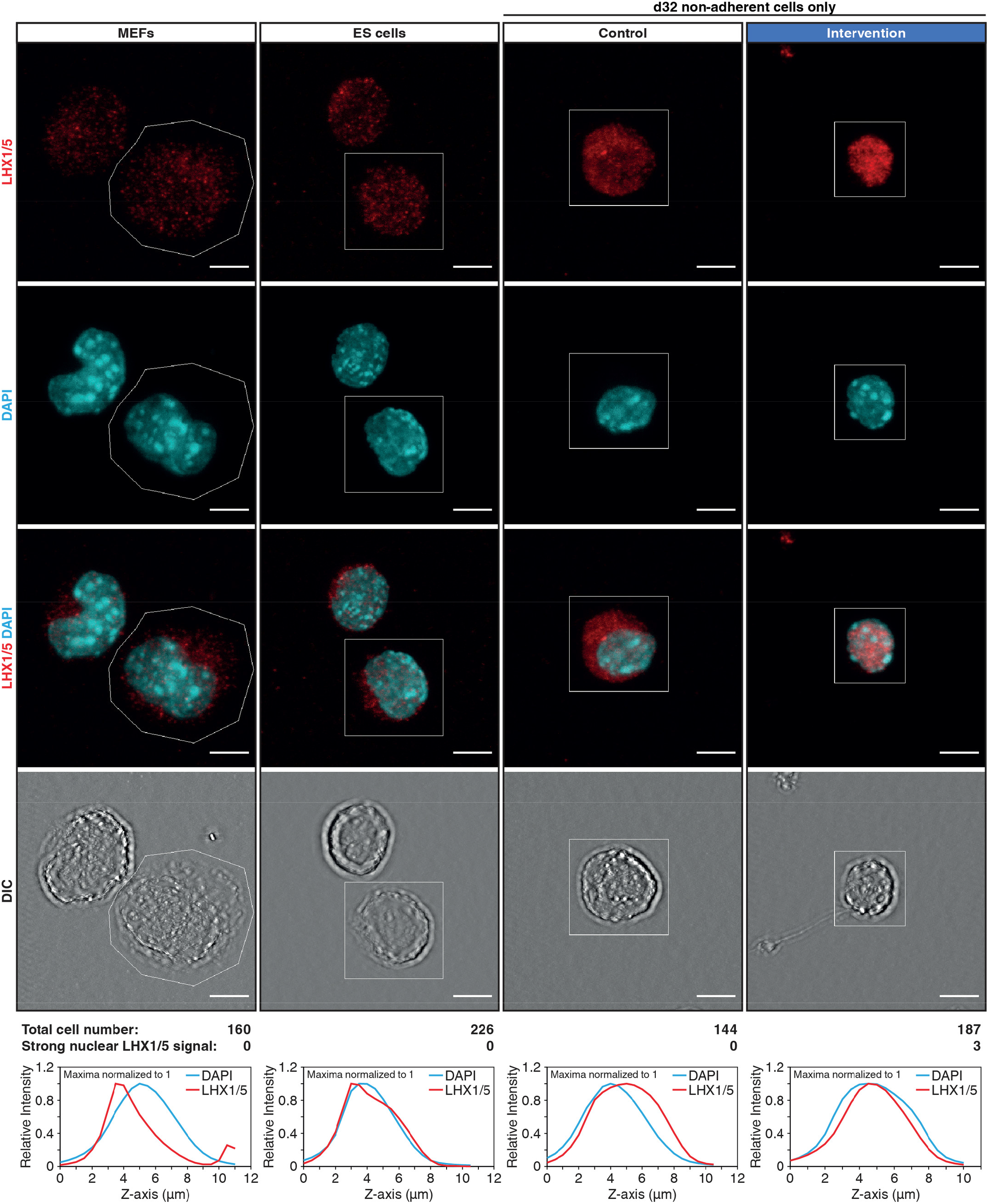
Zoom-in immunofluorescence images of LHX1/5. Zoom-in immunofluorescence images of LHX1/5 (red) in MEFs, ES cells, and CSMs in the control and intervention group, shown as maximum z-projections. DNA was counterstained with DAPI (blue). MEFs and ES cells were dissociated with accutase, while CSMs were obtained by collecting the non-adherent cell fraction of day 32 samples without enzymatic dissociation. After collection, all cells were stored alive for more than 6 months in liquid nitrogen before processing. DIC images at the bottom show the membrane boundaries of cells. The z-axis profiles (maximum intensities normalized to 1) of LHX1/5 and DAPI are plotted for regions marked by a white line in the respective images. LHX1/5 signal was detectable in the cytoplasm and nucleus of some cells in all samples. A strong nuclear LHX1/5 signal was detected in 3 out of 187 cells in CSMs in the intervention group but not in the other analyzed cells (MEFs: 160 cells, ES cells: 226 cells, CSMs in control group: 144 cells) (S3 and S4 Fig). Congruence of the LHX1/5 and DAPI signal in the images and z-axis profiles define a nuclear signal. Scale bar = 5 μm.

## Conclusions

Here, we demonstrate a new *in vitro* culture system using male mouse ES cells to study and manipulate aspects of spermatogonia development, including changes in gene expression and the generation of CSMs. We show that a transient chemical intervention at a time before upregulation of *Tex101* expression changes the differentiation process and leads to an enrichment of *Lhx1* expression in the resulting CSMs.

## Limitations

This is a cell culture-based *in vitro* differentiation system, where we used commercially available male mouse C57BL/6N ES cells (Millipore) that had a duplication of chromosome 8. It was previously reported that ES cells with trisomy 8 have a diminished efficiency to contribute to the germline of chimeric mice [14]. While our results showed that aspects of spermatogonia development, including changes in gene expression and cell morphology could be induced in these ES cells *in vitro*, further studies using euploid ES cells from different genetic backgrounds are necessary to prove that the observed results are not specific to the ES cells used here. We did not use mouse testis tissue or isolated SSCs as a comparison in this study. Therefore, it would be interesting to directly compare CSMs to SSCs, including gene expression analysis and studies of biological functionality in testis transplantation experiments.

## Materials and Methods

### Ethics statement

No animals were used in this study. All cells used in this study, consisting of the commercially available ES cell line purchased from Millipore (Millipore, SCC050) and the MEF line, did not require an ethics approval in accordance with the ETH Zurich Animal Welfare Office. All experiments were conducted according to the standard ethical guidelines of the ETH Zurich.

### Maintenance cell culture

All cells were cultured in an incubator with a humidified atmosphere at 37°C and 5% (v/v) CO_2_. Male (xy) mouse C57BL/6N ES cells (Millipore, SCC050) (S1 Fig) were cultured in T25 or T75 tissue culture flasks (TPP, Switzerland) that were coated with 0.1% gelatin solution (Millipore, ES-006-B). ES cells were passaged every 2 days with accutase (Thermo Fisher Scientific, A1110501). Basic medium (BM) consisted of Embryomax DMEM (Millipore, SLM-220-B), supplemented with 15% (v/v) Embryomax fetal bovine serum (FBS) (Millipore, ES-009-B), 1 mM sodium pyruvate (Millipore, TMS-005-C), 0.1 mM non-essential amino acids (Thermo Fisher Scientific, 11140050), and 2 mM Glutamax-I supplement (Thermo Fisher Scientific, 35050061). ES cells were cultured in 2iL medium, consisting of BM supplemented with 0.1 mM βME (Thermo Fisher Scientific, 31350010), 1000 units/ml LIF (Millipore, ESG1107), 1 µM CHIR99021 (Cayman Chemical, 13122) and 1 µM PD 0325901 (Cayman Chemical, 13034). Immortalized MEFs were kindly provided by Reinhard Fässler (Max Planck Institute of Biochemistry, Germany). MEFs were cultured in DMEM (Thermo Fisher Scientific, 21885) supplemented with 10% (v/v) FBS (Biowest, S181H).

### Intervention & control culture

For the intervention and control group the procedure illustrated in Fig 1 was followed. In the control group βME was always supplemented, while Ex-527 (Cayman Chemical, 10009798), RG-108 (Cayman Chemical, 13302) and tBHQ (Sigma-Aldrich, 112941) were replaced with dimethyl sulfoxide (DMSO) with the same final concentration as in the intervention group. Ex-527, RG-108 and tBHQ were added from stock solutions in DMSO. The intervention and control group were identical in all other aspects of the procedure. On day -24, ES cells were defrosted and cultured for 2 days in 2iL medium. On day -22, the medium of the intervention group was changed to 2iL medium supplemented with 5 µM Ex-527 and 100 µM RG-108 and the cells were passaged 1:5 to 1:10 every 2 days for 10 cell passages (20 days) in T25 or T75 tissue culture flasks. On day -2, cells were seeded into gelatin-coated 6-well plates (9.5 cm^2^ growth area per well, TPP, Switzerland) at 2×10^4^ cells/cm^2^ (3 ml/well) and cultured for 2 days in the same medium. No further cell passaging was performed from this point onwards. The total DMSO concentration was 0.03% (v/v). On day 0, the medium of the intervention group was changed to BM supplemented with 5 µM Ex-527, 100 µM RG-108 and 10 µM tBHQ and the total volume was increased to 6 ml/well. βME and 2iL were not present. The cells were cultured in this medium for 7 days with partial medium replacement of 3 ml/well (half of well volume) two times per day at hour 8 and hour 16. The total DMSO concentration was 0.04% (v/v). On day 7, the medium of the intervention group was changed to BM supplemented with βME, while keeping the same replacement frequency, volume and total volume. Until day 19, there was no noticeable cell loss during medium replacements because all cells were strongly adherent. From day 19 until day 32, the partial medium replacements were changed to 2 ml/well (one third of well volume) pipetting slowly from the edge of the well with a 1000 µl pipette to avoid the loss of non-adherent CSMs, while keeping the same replacement frequency, medium composition and total volume.

### Collection of cells

For day 0, day 7 and day 19 samples all cells were adherent and collected using 2 ml/well accutase (Thermo Fisher Scientific, A1110501) and a cell scraper, followed by centrifugation at 300 g for 5 minutes and washing with 1 ml phosphate-buffered saline (PBS, pH 7.4). For day 32 samples, only the non-adherent cells were collected by rinsing the well 20-times with its supernatant using a 5 ml pipette, followed by filtration of the supernatant through a 100 µm cell strainer (Corning, 352360). Part of the non-adherent cells was centrifuged at 300 g for 5 minutes, washed with 1 ml PBS and immediately used for RNA isolation, while the rest was frozen in cryopreservation medium (CryoStor CS10, STEMCELL Technologies, 07930) in liquid nitrogen for later immunofluorescence.

### Real-time PCR analysis

Total RNA was isolated from ∼0.5 to 2×10^6^ cells per sample using the RNeasy Plus Mini Kit (Qiagen, 74134) with RNase-free DNase treatment (Qiagen, 79254) according to the manufacturer’s protocol. For reverse transcription (RT) 2 μg total RNA was reverse transcribed into cDNA in a reaction volume of 20 μl using the iScript™ Advanced cDNA Synthesis Kit (Bio-Rad, 1725038). Real-time PCR reactions were performed on the resulting cDNA using the CFX Connect™ Real-Time PCR Detection System (Bio-Rad, USA) and SsoAdvanced™ Universal SYBR^®^ Green Supermix (Bio-Rad, 1725271). PCR reactions were incubated at 95°C for 3 minutes, followed by 39 cycles of 95°C for 10 s and 62°C for 30 s; and were run as five replicates pooled from three independent experiments. PCR Primers were designed using PrimerSelect from the Lasergene software package (DNASTAR, USA). All primers were synthesized at Microsynth. Results were analyzed using CFX Manager™ software (Bio-Rad, USA) and relative gene expression levels were calculated based on the ΔΔCt method with normalization using *Gapdh*. Gene abbreviations, NCBI reference sequences for mRNA (used as template for primer design), amplicon size and primer sequences are indicated in S1 Table.

### Statistical analysis

Statistically significant differences between data values were calculated with Prism 8 (GraphPad, USA) using one-way ANOVA and the Tukey post hoc test (one star represents p < 0.05, two stars represent p < 0.01, three stars represent p < 0.001 and four stars represent p < 0.0001).

### Immunofluorescence

100 µl of cells previously frozen in cryopreservation medium in liquid nitrogen (0.5*10^6^ cells) were defrosted and transferred to Sterilin™ Clear Microtiter™ Plates (Thermo Fisher Scientific, 611V96) prefilled with 200 µl PBS and centrifuged at 300 g for 5 minutes. To avoid high cell loss, all liquid removals in the microtiter wells were performed carefully from the wall of the well by capillary action with the pointy end of twisted Kimtech Science™ Precision Wipes (Kimberly-Clark Professional, 05511) that were fitted into wide bore 1000 µl pipette tips. Cells were washed three times with 250 µl PBS and centrifugation at 300 g for 5 minutes. Cells were resuspended in 30 µl PBS, followed by addition of 250 µl formaldehyde (2.5% w/v in PBS) and fixed for 10 minutes at room temperature. Cells were centrifuged at 500 g for 5 minutes and the formaldehyde removed. Cells were washed once with 250 µl PBS and centrifugation at 500 g for 5 minutes. Cells were then resuspended in 500 µl PBS, transferred to a 2 ml microcentrifuge tube and stored at 4°C.

A tailored immunofluorescence protocol was used to achieve a high local cell density, while minimizing cell loss. Fixed cells were adhered to Superfrost™ Plus Gold Adhesion Slides (Thermo Fisher Scientific, K5800AMNT72) which showed the best adhesion across fixed CSMs, ES cells and MEFs, compared to several other commercial microscope slides. A sandwich of two silicone sheetings (cut to 15 mm x 15 mm, 0.010 inches NRV G/G 40D, SMI Specialty Manufacturing) with differently sized concentric holes defined the cell adhesion and washing areas. A small hole (4 mm diameter) in the bottom sheeting defined the cell adhesion area on the slide (∼0.1257 cm^2^) and a large hole (9 mm diameter) in the top sheeting defined the washing area (S5A Fig). This setup reduced shear stress in the adhesion area by keeping it liquid-filled during the washes. During all incubations, the slide was placed in a container together with a separate dish of water to minimize evaporation. 20 µl of fixed cells (7×10^4^ cells/ml) were placed on the adhesion area (∼1.1×10^4^ cells/cm^2^) and incubated for 1 hour at room temperature. Cells were permeabilized with 0.3% Triton X-100 in PBS for 10 minutes, blocked with 5% goat serum in PBS (Sigma-Aldrich, G9023) for 1 hour at room temperature, and stained using a standard immunofluorescence protocol. Antibody incubations were performed in 5% goat serum in PBS for 1 hour at room temperature for primary and secondary antibodies. Nuclei of cells were stained with 5 µg/ml DAPI in PBS for 10 minutes at room temperature. Primary antibodies were mouse anti-LHX1/5 (monoclonal, 5 µg/ml, Developmental Studies Hybridoma Bank, 4F2 supernatant, antigen used to raise antibody: Lim 1+2 / LhxV5, antigen species: rat). Secondary Antibodies were goat anti-mouse IgG H&L Alexa Fluor 647 (polyclonal, 1:200, Abcam, ab150115). After staining, cells were washed three times with ultrapure water to avoid salt crystals. The liquid in the adhesion area was then carefully removed with a twisted Kimtech Science™ Precision Wipe without disturbing the stained cells, followed by removal of the sandwiched silicone sheeting. A cover glass (0.17 mm diameter, 18 mm x 18 mm, Carl Zeiss, 10474379) was placed on the adhesion area and fixed with scotch tape at the top edge. 20 µl of ProLong™ Gold Antifade Mountant (Thermo Fisher Scientific, P10144) pipetted to the lower edge of the cover glass filled the space by capillary force (S5B Fig). Cells were mounted overnight at room temperature, followed by sealing of the cover glass with CoverGrip™ Coverslip Sealant (Biotium, 23005) and storage at 4°C. 12-bit numerical images were acquired as z-Stacks (0.5 µm step size) with a Leica TCS SP8 confocal microscope (Leica Microsystems, Germany) with a HC PL APO CS2 63x/1.40 oil objective. The pinhole was set to 1.0 airy unit. The resolution was set to 512 pixels x 512 pixels with a field of view of 246 µm x 246 µm for zoom-out images (pixel size = 481 nm) (S3 Fig) and 30.75 µm x 30.75 µm for zoom-in images (pixel size = 60 nm) (Fig 4 and S4 Fig). Alexa Fluor 647 was excited with a 633 nm laser and DAPI was excited with a 405 nm laser. Images were processed as maximum z-projections using the ImageJ software (http://rsbweb.nih.gov/ij/). Differential interference contrast (DIC) images were further processed with a fast Fourier transform (FFT) shortpass filter filtering large structures down to 15 pixels, with additional suppression of horizontal stripes with a tolerance of direction of 95% for DIC zoom-in images, and adjustment of brightness/contrast using the ImageJ software.

### Light microscopy

The live cells were imaged on day 0, day 7, day 19 and day 32 using an Axiovert 200 M microscope (Carl Zeiss, Germany) with a LD Plan-Neofluar 20x/0.4 Korr Ph2 objective.

### Cytogenetic analysis by FISH

For chromosome painting by fluorescence in situ hybridization (FISH), ES cells were cultured in 2iL medium on gelatin-coated 6-well plates to 80% confluency. Cells were arrested in metaphase using 0.5 ml Colcemid (10 mg/ml stock solution) per 10 ml medium for 2 hours. After delicate centrifugation (120 g, 8 minutes, 24°C) and removing the supernatant, hypotonic KCl solution (0.075 M, 37°C) was added by slow dropping up to 6 ml. Cells were then incubated for 20 to 25 minutes at 37°C. After incubation, centrifugation and removing the supernatant, cells were fixed by slow dropping of frozen (−30 to -80°C) Carnoy fixative (absolute methanol and glacial acetic acid in a 3:1 ratio) up to 8 ml, centrifugation (120 g, 8 minutes, 4°C), and after removing the supernatant adding again fixative up to 4 ml. The chromosome suspensions were concentrated to 100 μl of which 10 μl were dropped onto cold, wet microscope slides. All materials dropped on the slides covered a circular area of less than 15 mm diameter. The microscopic preparations were then incubated in a dry environment at 37°C for 24 hours until hybridization. Hybridizations were carried out for 18 to 24 hours. In a first step, RNA was digested by treatment with 100 to 120 μl of RNase A in 2x saline-sodium citrate buffer (SSCB, 100 µg/ml) in a humidified atmosphere for 1 hour at 37°C. Slides were then rinsed three times for 2.5 minutes in 2x SSCB at room temperature, and then dehydrated by incubation in a graded series of ethanol (70, 80, and 90%) for 2.5 minutes each at room temperature. Multicolor FISH was performed for X and Y chromosomes using mouse paint chr.X red and mouse paint chr.Y aqua probes (ChromBios, Germany, PMXRD and PMYAQ, respectively) and for all chromosomes using the 21XMouse Chromosome Paint Kit (MetaSystems, Germany, D-0425-060-DI) according to the manufacturer’s protocol. Images were acquired at 1200x magnification with an Axio Imager Z1 microscope (Carl Zeiss, Germany) using the MetaSystems software ISIS/IKAROS (MetaSystems, Germany).

## Acknowledgments

We thank Reinhard Fässler (Max Planck Institute of Biochemistry, Germany) for kindly providing us with mouse embryonic fibroblasts, and Justine Kusch-Wieser for help with confocal microscopy (ScopeM ETH Zurich, Switzerland).

## Author Contributions

**Cameron Moshfegh:** Conceptualization, Data curation, Formal analysis, Investigation, Methodology, Data visualization, Validation, Software, Project administration, Supervision, Funding acquisition, Original draft, Review & editing. **Sebastian Giovanni Rambow:** Investigation, Methodology, Review & editing. **Seraina Andrea Domenig:** Investigation, Methodology, Review & editing. **Aldona Pieńkowska-Schelling:** Investigation, Data curation, Review & editing. **Ulrich Bleul:** Supervision, Review & editing. **Viola Vogel:** Resources, Supervision, Review & editing.

## Supplementary Figures and Tables

**S1 Fig.**
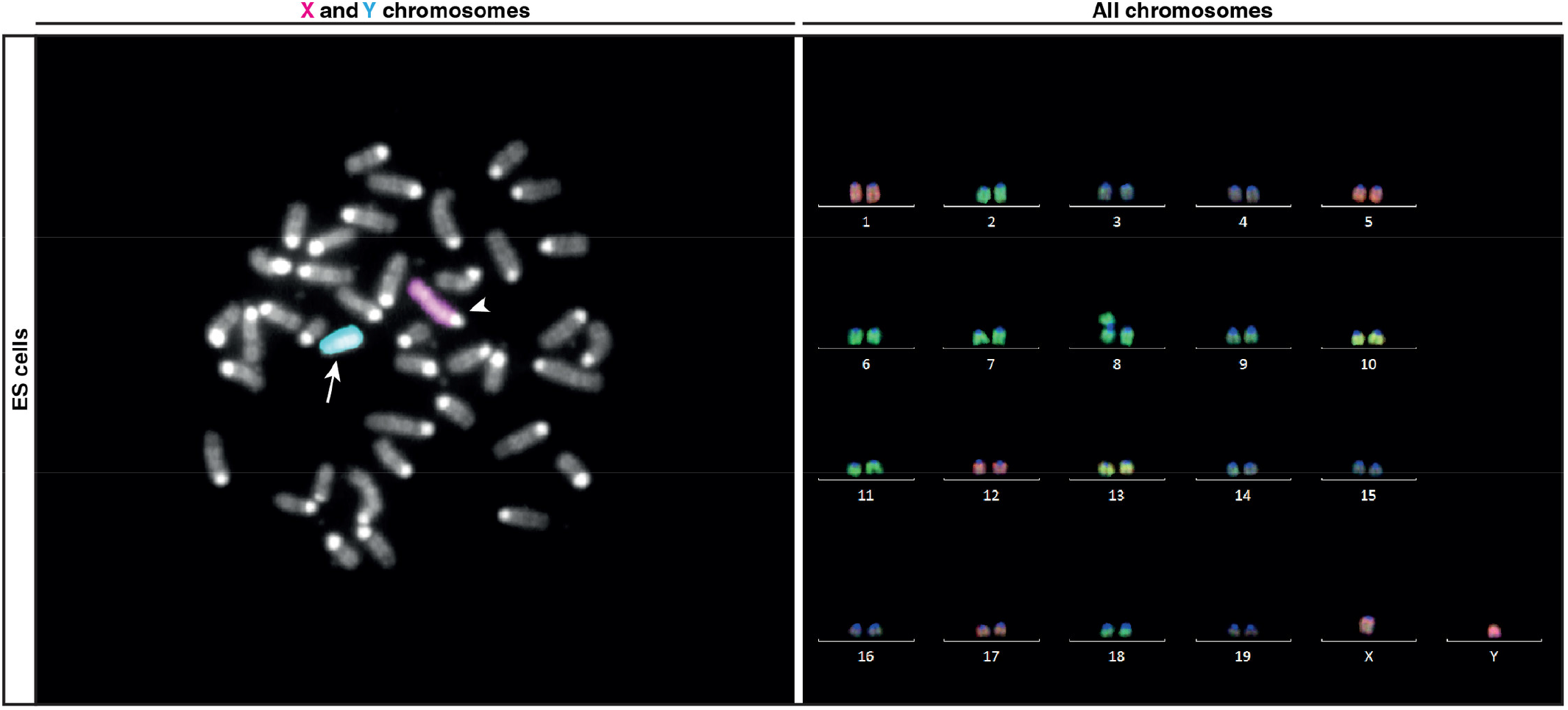
Multicolor FISH of metaphase chromosomes in ES cells. Sex chromosome analysis of 4 metaphase spreads all showed a male genetic sex. X (red, arrowhead) and Y (blue, arrow) chromosomes confirm a male genetic sex of the ES cells (left image). Complete karyogram analysis of 3 metaphase spreads all showed a duplication of chromosome 8 with chromosome counts ranging from 40 to 42. The complete karyogram shows an abnormal metaphase with 41 chromosomes with a trisomy in chromosome 8. Sorted autosomes (chromosomes 1 to 19) and sex chromosomes (chromosomes X and Y) are shown (right image).

**S2 Fig.**
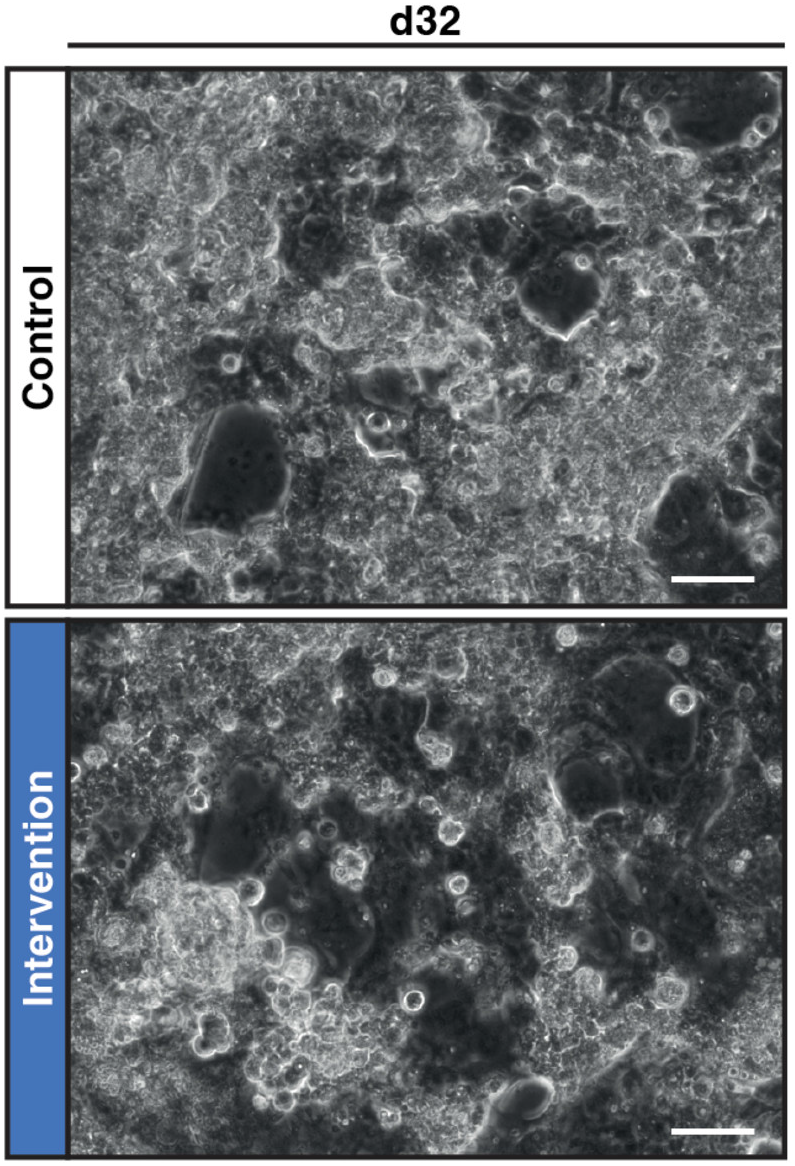
Differential progress of cell dissociation in the control and intervention group. Phase contrast images of the differentiation process of ES cells to CSMs in the control (white) and intervention group (blue) on day 32, showing a more progressive dissociation of cell sheets in the intervention group compared to the control group. Scale bar = 50 μm.

**S3 Fig.**
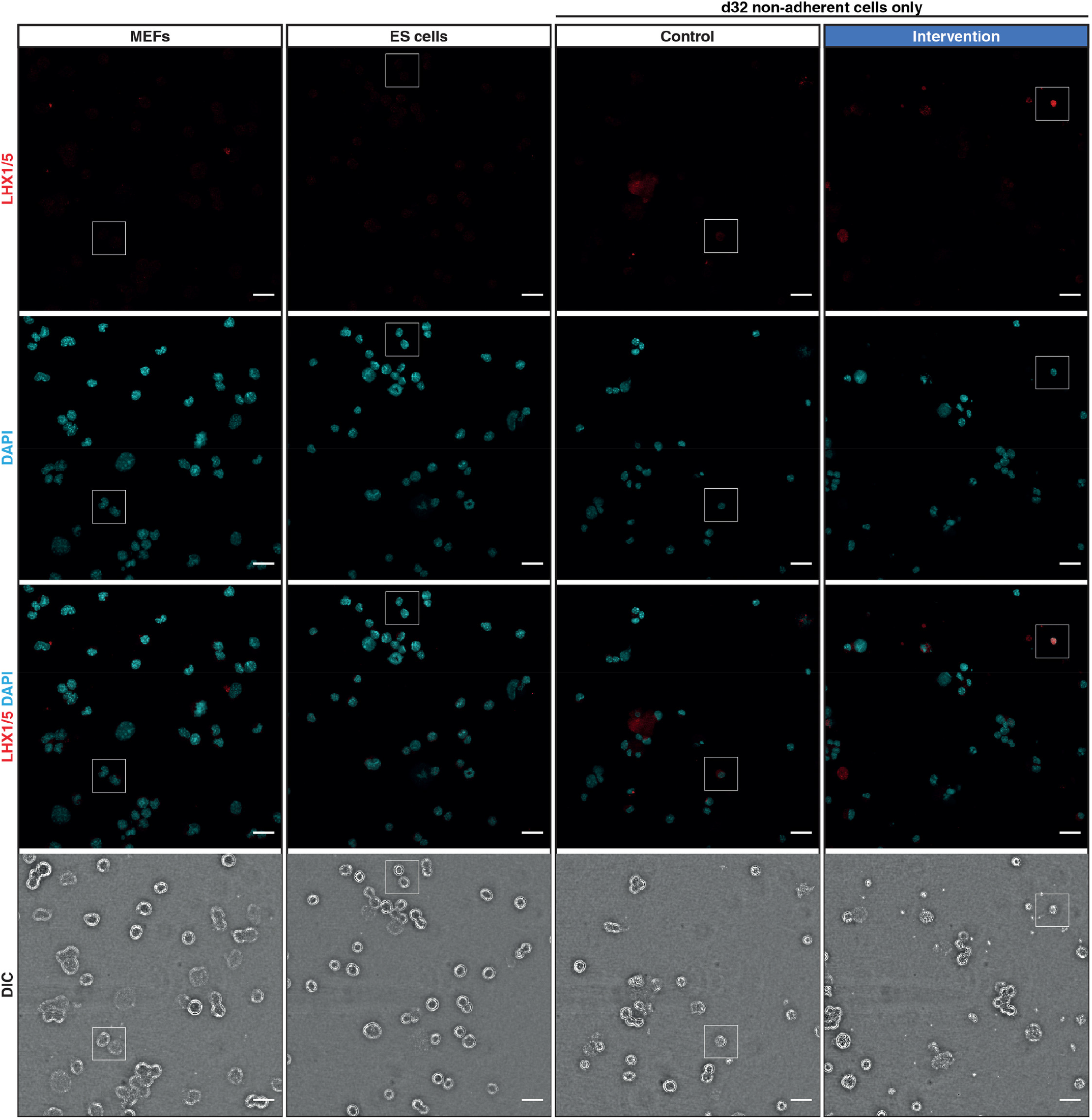
Zoom-out immunofluorescence images of LHX1/5. Zoom-out immunofluorescence images of LHX1/5 (red) in MEFs, ES cells, and CSMs in the control and intervention group, shown as maximum z-projections. The regions marked by a white line correspond to the respective zoom-in images in Fig 4. DNA was counterstained with DAPI (blue). MEFs and ES cells were dissociated with accutase, while CSMs were obtained by collecting the non-adherent cell fraction of day 32 samples without enzymatic dissociation. After collection, all cells were stored alive for more than 6 months in liquid nitrogen before processing. DIC images at the bottom show the membrane boundaries of cells. Scale bar = 20 μm.

**S4 Fig.**
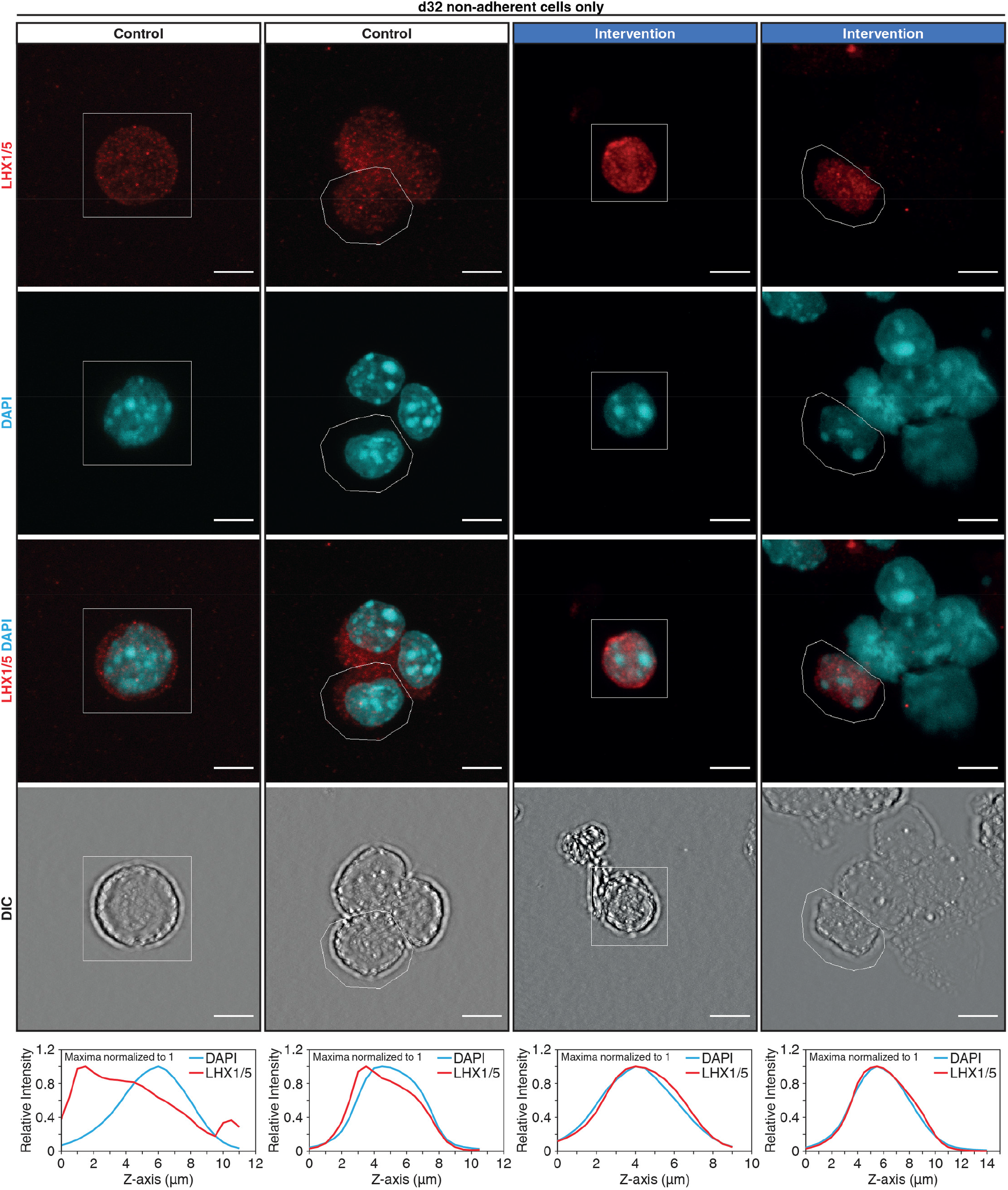
Additional zoom-in immunofluorescence images of LHX1/5. Additional zoom-in immunofluorescence images of LHX1/5 (red) in CSMs in the control and intervention group, shown as maximum z-projections. The other two CSMs in the intervention group with a strong nuclear LHX1/5 signal are shown. DNA was counterstained with DAPI (blue). CSMs were obtained by collecting the non-adherent cell fraction of day 32 samples without enzymatic dissociation. After collection, all cells were stored alive for more than 6 months in liquid nitrogen before processing. DIC images at the bottom show the membrane boundaries of cells. The z-axis profiles (maximum intensities normalized to 1) of LHX1/5 and DAPI are plotted for regions marked by a white line in the respective images. Congruence of the LHX1/5 and DAPI signal in the images and z-axis profiles define a nuclear signal. Scale bar = 5 μm.

**S5 Fig.**
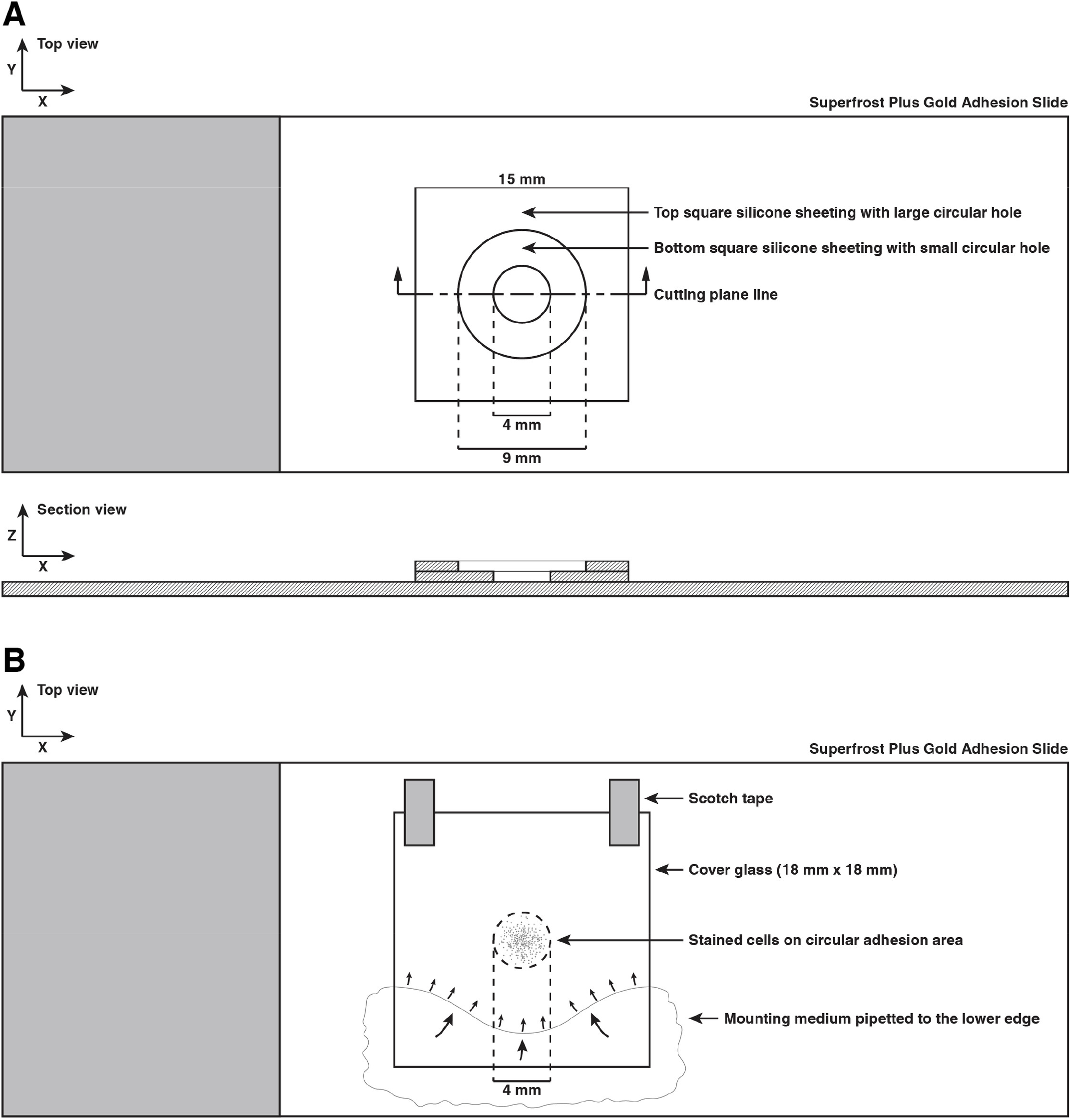
Schematics for the immunofluorescence staining setup. (A) A schematic representation for the immunofluorescence staining of in-solution fixed cells. A sandwich of two silicone sheetings with differently sized concentric holes defined the cell adhesion and washing areas. The xy top view and the xz section view are shown. See Materials and Methods for a detailed description. (B) A schematic representation for the mounting of stained cells. Mounting medium was pipetted to the lower edge of a cover glass that was fixed with scotch tape on top of the stained cells and filled the space by capillary force. The xy top view is shown. See Materials and Methods for a detailed description.

**S1 Table.**
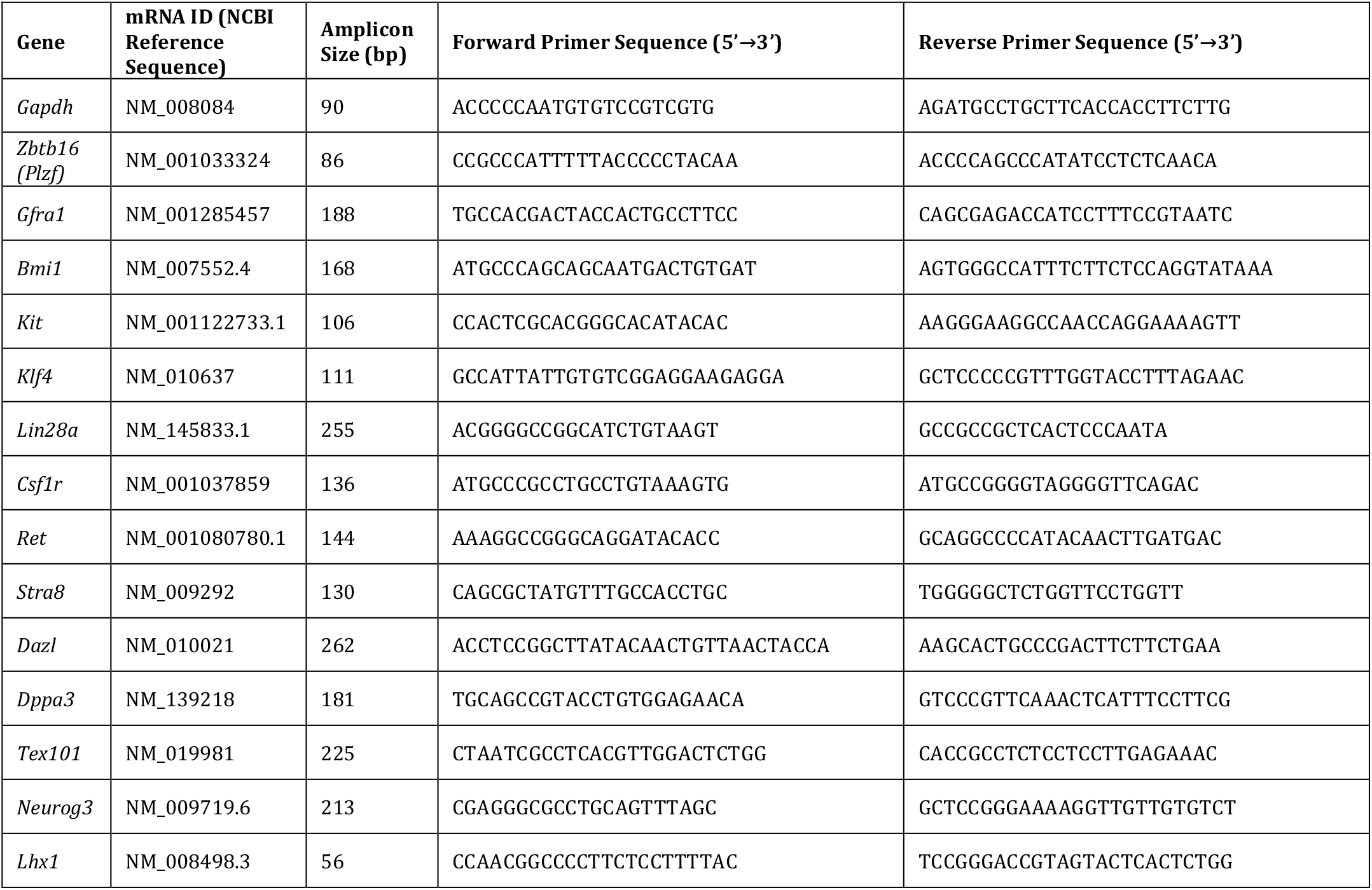
Primer sequences. Gene abbreviations, NCBI reference sequences of mRNAs (used as template for primer design), amplicon size and primer sequences for the real-time PCR analysis (See Materials and Methods for a detailed description).

